# Endogenous Retrovirus Loci and Induced Changes in Gene Expression in Japanese Indigenous Chickens

**DOI:** 10.1101/2024.08.05.606564

**Authors:** Shinya Ishihara, Jun-ichi Shiraishi, Saki Shimamoto, Daichi Ijiri

**Author notes:** Corresponding author: Shinya Ishihara, Address: Department of Animal Science, Nippon Veterinary and Life Science University, 1-7-1 Kyonancho, Musashino, Tokyo, Japan, 180-8602.

## Abstract

When retroviruses infect germ cells and are transmitted to offspring, they become endogenous retroviruses (ERVs), whose insertions influence the expression of nearby genes. This study aimed to identify the genomic loci of ERVs in commercial broiler (Ross308), Tosa-Jidori, and Yakido chickens as well as to elucidate their impact on neighboring gene expression. Whole-genome data were obtained using next-generation sequencing, and candidate ERV loci were identified using the RetroSeq software. The Integrative Genomics Viewer tool was used to confirm target site duplications (TSDs) as evidence of ERV insertions. All reads within 200 bp of these TSDs were extracted to create contigs, confirming the presence of ERV sequences in the contigs using BLASTN. Gene expression levels were estimated by focusing on genes located near the 172 identified ERV loci. Among these, 119 loci were detected in broiler chickens, 80 in Tosa-Jidori chickens, and 86 in Yakido chickens, with 28 loci shared among them. Moreover, of these 172 loci, 75 were located within or near genes. Significant differences in gene expression were observed for N-acetylated alpha-linked acidic dipeptidase 2, glypican 6, and phospholipid scramblase family member 5 depending on the presence of ERV insertions. These results suggest that ERV insertions may influence the expression of certain genes, providing insights into the genetic diversity and evolutionary background of commercial and indigenous chickens. Understanding the effects of ERV insertions on gene expression can inform future genetic research and poultry breeding programs aimed at improving health and productivity.

## Background

Endogenous retroviruses (ERVs) are ancient genetic elements comprising virus-derived sequences that have been integrated into the genomes of various organisms. These are formed when viruses insert themselves into a host genome and are subsequently transmitted to the its progeny. Structurally, ERVs possess long terminal repeats (LTRs) at both ends and encode structural proteins (*gag*), reverse transcriptases (*pol*), and envelope proteins (*env*). These sequences accumulate mutations and become fragmented over generations, often leading to a loss of function in many ERVs. Once integrated into the genome, ERVs can influence the transcription and expression of nearby host genes, primarily through regulatory elements present in their LTR sequences, such as promoters and enhancers [1, 2, 3]. Recently, ERVs have gained significant attention as valuable markers for understanding genetic relationships and evolutionary processes among species. Because they are inherited vertically across generations, the presence or absence of ERVs in different species may elucidate phylogenetic relationships. While shared ERVs among different species indicate a common ancestral infection, species-specific ERVs suggest independent integration events [4]. Furthermore, as ERVs are genetically stable over long evolutionary timescales, they make reliable markers. Unlike other transposable elements, most ERVs cannot retrotranspose nor spread within the genome, resulting in relatively stable genomic distributions. Consequently, the evolutionary history of a species can be traced by analyzing its ERV integration patterns [5, 6]. In avian species, researchers have focused on the endogenous avian retrovirus (EAV) family and avian leukosis virus subgroup E (ALVE) [7, 8, 9, 10, 11, 12], reporting that the blue eggshells of Araucana chickens are influenced by ERVs. Understanding the role of ERVs in avian species provides significant insight into their evolutionary biology and genetic diversity [13].

Japanese indigenous chickens comprise approximately 50 breeds, most of which are classified as fancy or practical breeds based on traits such as unique crowing, ornamental feathers, and fighting characteristics, as well as egg and meat production [14, 15, 16, 17, 18]. Examples of these breeds include Tosa-Jidori, the oldest type of Japanese native chicken with a history of over 2000 years, and Yakido, a Malay-type breed with great height and black feathers [19, 20, 21]. A study using 20 microsatellite DNA markers to construct phylogenetic trees showed that these native Japanese breeds possess distinct genetic backgrounds compared to those of foreign breeds such as White Leghorn and Rhode Island Red [22]. These diverse Japanese indigenous chicken breeds, which underwent unique evolutionary processes under different environmental and artificial selection pressures, serve as intriguing models for studying the insertion sites and effects of ERVs. For instance, breeds that are adapted to harsh climates or specific local conditions may exhibit unique ERV profiles.

This study aimed to elucidate how ERVs contribute to the genetic diversity and adaptation of Japanese indigenous chickens, which may provide insights for breeding programs and conservation strategies. To achieve this objective, we identified the loci of ERVs in the genomes of commercial broiler (Ross308), Tosa-Jidori, and Yakido chickens and elucidated how these affect gene expression. We employed whole-genome sequencing technologies and bioinformatic analyses to identify ERV insertion sites and assess their regulatory impacts. Our findings suggest that ERV insertions may affect the expression of neighboring genes and demonstrate their contribution to the genetic diversity of these chickens.

## Results

### Sequencing data quality and identification of the non-reference ERV breakpoint

In this study, 150-bp paired-end reads were aligned using the Burrows-Wheeler Aligner and Mem (BWA-mem) algorithm [23], achieving an overall mean sequence depth of 44.6x (ranging 38.6–57.5) across all chicken samples (Table S1); the detailed mapping results are presented in Table S1. Over 99% of the paired-end reads for each chicken were successfully aligned with the reference Gallus genome, while 1.72–2.77% were classified as improperly mapped reads. Additionally, 0.13–0.16% of the reads were singletons, aligning to a single side. The RetroSeq software [24] identified 167–240 candidate ERV insertion loci per individual (Table S1). In total, 184 ERV loci were identified across the three types of chicken. Of the 184 detected ERV loci, 10 matched the surrounding sequences of the reference genome, and 2 were shorter than 200 bp; consequently, these were excluded from further analysis. The contigs derived from the remaining 172 loci were used in the analysis (Table S2). The distribution of the 172 loci within the genome was as follows: 57 on Chr1, 33 on Chr2, 23 on Chr3, 19 on Chr4, and 12 on ChrZ, representing 81.4% of the total. A strong correlation was observed between chromosome length and the number of detected loci (Pearson’s correlation coefficient = 0.96). Broiler, Tosa-Jidori, and Yakido chickens had 117, 78, and 82 ERV loci, respectively. The number of loci shared among the breeds is shown in Figure 1, with 28 loci shared among all types of chicken. The ERV loci specific to broiler, Tosa-Jidori, and Yakido chickens were 54, 18, and 23, respectively. Of the 172 loci, 95 were shared with junglefowls, 94 of which were shared with red junglefowls, the ancestor of domestic chickens [25, 26], while the remaining locus was shared with gray junglefowls (Figure S1). The summary statistics for heterozygosity in each breed are presented in Table 1. Broilers had higher estimates of observed heterozygosity (Ho=0.28) and expected heterozygosity (He=0.23) than those of Tosa-Jidori (Ho=0.17, He=0.14) and Yakido (Ho=0.19, He=0.15) chickens. In the constructed phylogenetic tree (Figure 1), ERVs were frequently observed at the same loci in the same chicken, although these were not consistently shared at identical loci (Table S2). For example, the number of homozygous ERV loci was 0 in broilers, 10 in Tosa-Jidori, and 14 in Yakido. Nonetheless, clusters in the phylogenetic tree were clearly delineated for chickens and junglefowls (Figure 1).

**Figure 1.**
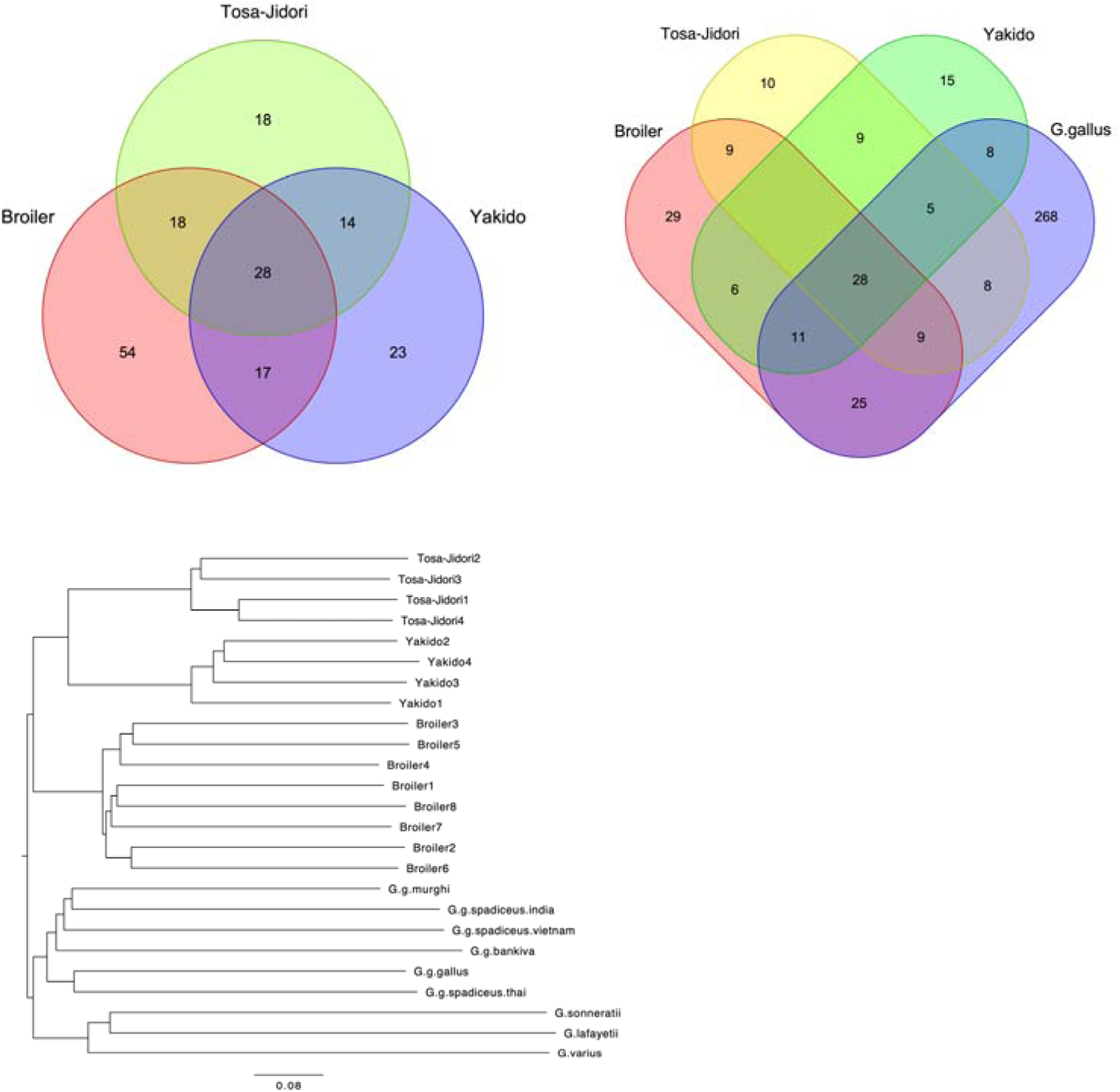
Number of endogenous retrovirus (ERV) loci detected among broiler, Tosa-Jidori, Yakido and gallus genus as well as phylogenetic trees. (A) Number of ERV loci across broiler, Tosa-Jidori, and Yakido and the overlap between each ERV locus. (B) Number of ERV loci across broiler, Tosa-Jidori, Yakido, and red junglefowl and the overlap between each ERV locus. (C) Phylogenetic tree constructed based on the presence or absence of ERV loci. The scale bar indicates the distance calculated from the presence/absence matrix.

**Table 1.** Comparison of Observed and Expected Heterozygosity in Different Chickens.

| Chicken | Observed heterozygosity | Expected heterozygosity |
| --- | --- | --- |
| Broiler | 0.28 | 0.23 |
| Tosa-Jidori | 0.17 | 0.14 |
| Yakido | 0.19 | 0.15 |

### Analysis of Detected ERV Types and Sequences

The contigs detected from the 172 loci were mapped to ERV sequences in the database (Table S3). The Chr1_159876144 locus was mapped to two locations in EAV-HP (previously described as ev/J, AF125529), while the Chr2_133314053 locus was mapped to two locations in the Murine leukemia virus (MLV) related retroviruse (DQ280312). The numbers of contigs mapped to EAV-HP, EAV-51, EAV-0, and EAV-0/E51 were 79, 9, 33, and 1, respectively (Table 2). Moreover, 8 and 1 contigs were mapped to ALVE and MLV, respectively. The lengths of the obtained contigs varied regardless of ERV type; however, the average length of contigs corresponding to ALVE (573.5 bp, 405–734 bp), was greater than that of other ERVs. Most of these contigs contained LTR sequences, with some including gag, pol, env, or gag/env fusion protein sequences (Figure S2).

**Table 2.** Contig length distribution of Various ERVs in Chicken Genomes.

| ERV | Accession No. | Mapped contigs | Contig length |  |  |  |  |  | Total |
| --- | --- | --- | --- | --- | --- | --- | --- | --- | --- |
|  |  |  | 201-30 | 301-40 | 401-50 | 501-60 | 601-70 | 701-80 |  |
|  |  |  | 0 | 0 | 0 | 0 | 0 | 0 |  |
| EAV-HP | AF125529 | 24 | 2 | 14 | 4 | 2 | 2 | 0 | 79 |
|  | AJ623291 | 47 | 36 | 6 | 2 | 2 | 1 | 0 |  |
|  | NC_005947.1 | 18 | 0 | 9 | 2 | 2 | 5 | 0 |  |
| EAV-51 | AM418553 | 9 | 6 | 2 | 1 | 0 | 0 | 0 | 9 |
| EAV-0 | AM418554 | 33 | 1 | 5 | 21 | 5 | 1 | 0 | 33 |
| EAV-0/E51 | AM418555 | 1 | 0 | 0 | 0 | 0 | 1 | 0 | 1 |
| ALVE | MT263508 | 8 | 0 | 0 | 1 | 4 | 2 | 1 | 8 |
| MLV | DQ280312 | 1 | 1 | 0 | 0 | 0 | 0 | 0 | 1 |
| LTR | M31063 | 9 | 6 | 3 | 0 | 0 | 0 | 0 | 31 |
|  | M31065 | 3 | 1 | 1 | 1 | 0 | 0 | 0 |  |
|  | M31066 | 19 | 13 | 1 | 5 | 0 | 0 | 0 |  |
| Total |  | 172 | 66 | 41 | 37 | 15 | 12 | 1 | 172 |

## Impact of ERV Insertions on mRNA Expression

The RNA expression levels, classified based on the presence or absence of ERV insertions at each locus, are shown in Figure S3. Of the loci with detected ERVs, 75 were located near or within genes (Table S2). To examine the relationship between the presence of these gene insertions and RNA expression, transcripts per million (TPM) values were calculated (Table S4). Among the 75 genes, 24 were excluded from the analysis for one of three reasons: all individuals had ERV insertions, only one individual had an insertion, or over 90% of the individuals showed no gene expression. Figure S3 shows how the presence or absence of ERV insertions for the remaining 51 genes influenced gene expression. Significant differences were observed in the expression of N-acetylated alpha-linked acidic dipeptidase 2 (*NAALAD2*), glypican 6 (*GPC6*), and phospholipid scramblase family member 5 (*PLSCR5*) (adjusted p-values = 0.0022, 0.0119, and 0.0119, respectively). In *NAALAD2* and *GPC6*, ERV insertions were associated with decreased expression, whereas in *PLSCR5*, they were associated with increased expression (Figure 2).

**Figure 2.**
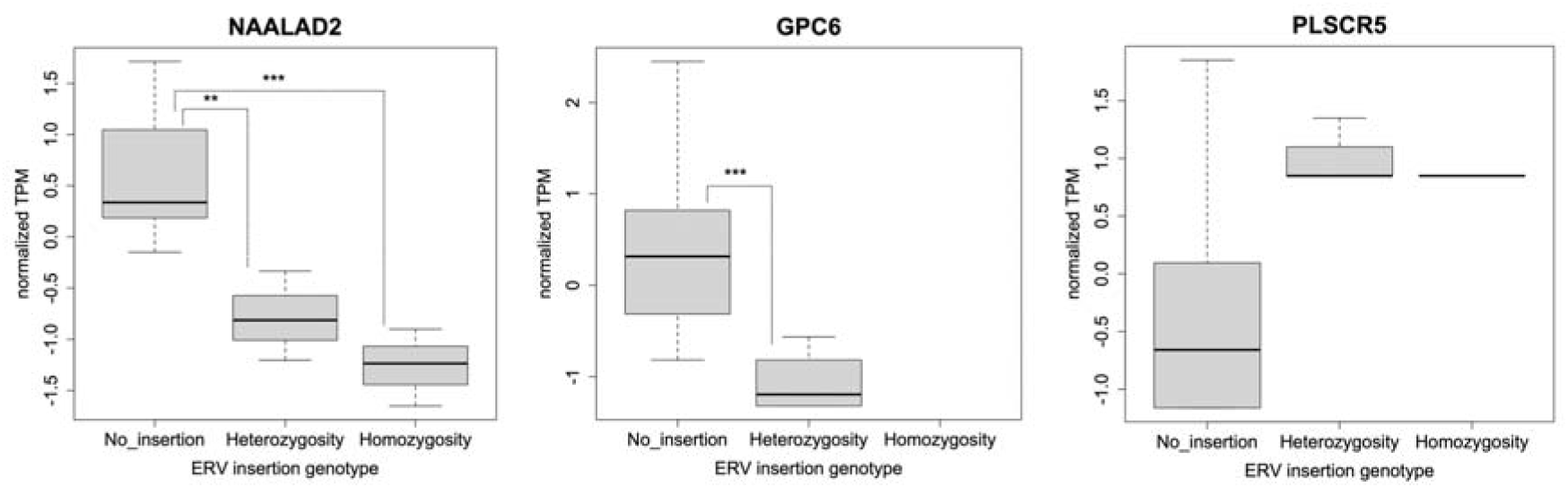
Normalized transcripts per million TPM) values for PLSCR5, NAALAD2, and GPC6. Box plots represent the distribution of gene expression levels across three ERV insertion genotypes: no insertion, heterozygosity, and homozygosity. Each box plot shows the median (line within the box), interquartile range (the box), and range of the data (whiskers). Statistical significance was determined using the Tukey honestly significant difference test, indicated by asterisks (adjusted p-value; *p<0.05, **p<0.01, ***p<0.001).

## Discussion

In this study, we employed next-generation sequencing to identify ERVs that were absent in a reference genome, rigorously trimming the obtained sequence data using stringent criteria. As more than 99% of all read pairs mapped to the Gallus reference genome, the assembled sequence data were considered to be high quality. We subsequently attempted to detect ERVs in broilers and Japanese indigenous chickens using improper pairs and singleton sequence reads that were incorrectly mapped to the reference genome. We identified 172 ERV loci across the three chicken types. Using a similar method, analysis of the genus Gallus—specifically, the red junglefowl—revealed the presence of 116 ERV loci (range: 61–123), which is largely consistent with previous findings [10]. In our study, the numbers of ERV loci detected in broilers, Tosa-Jidori, and Yakido chickens were 119, 80, and 86, respectively, which are comparable to previous findings. Additionally, we visually confirmed the breakpoints detected by RetroSeq using Integrated Genome Viewer (IGV) to identify target site duplications (TSDs) and included the ERV sequences in the contigs assembled from the surrounding sequences. Our results demonstrate the validity of the methods used for detecting ERVs in different chicken breeds and species, confirming their high reliability.

The ERV structures at each locus were inferred from the contigs. However, because of the tendency of ERVs to accumulate in repetitive regions [27, 28, 29], short-read sequencing alone may have limitations in accurately detecting complete ERV structures, including the *gag*, *pol*, and *env* regions, or solo-LTRs. To overcome these limitations, single-molecule real-time sequencing or nanopore sequencing may be used to determine complete ERV insertion sequences. This would allow for high-resolution mapping, precise localization of gene insertions, and the identification of mutations. Combining short- and long-read sequencing is expected to improve the differentiation of heterozygosity and homozygosity, leading to more accurate localization of gene insertions and mutations as well as a more refined functional analysis of genetic variation.

In this study, we analyzed the ERV insertion sites in the chicken genome to elucidate the genetic diversity and evolutionary background of different chickens. The strong correlation between chromosome length and the number of insertion sites suggests that ERV insertions occur randomly throughout the genome. However, as we did not account for existing ERVs in the reference genome, determining the concentration of ERV insertions in specific genomic regions conclusively was not possible. We found that broilers exhibited higher heterozygosity than that of other chickens, indicating greater ERV diversity. Our phylogenetic analysis based on the presence or absence of ERVs revealed distinct clusters for each chicken, with ERVs frequently observed at the same insertion sites within the same chicken type. This suggests that the observed ERV loci are fixed within specific lineages through selection pressure or genetic drift during breed formation. However, instances of ERV locus loss owing to genetic drift were also observed. For example, while the murine leukemia virus-related ERV (DQ280312) was confirmed in all junglefowls, it was not detected in two broilers. The fixation or loss of ERVs is influenced by selection pressure and genetic drift within the host population [30]; studies on commercial chickens have also reported that certain ERV insertion sites are fixed owing to selection pressure. The findings obtained here suggest that rare indigenous chickens possess unique ERV loci. The high heterozygosity of ERVs in broilers indicates a possible crossbreeding with populations possessing diverse ERV loci during breed formation. Of the 172 detected ERV loci, 94 matched those reported in red junglefowl, while the phylogenetic analysis placed commercial and indigenous chicken subspecies in clusters close to that of the red junglefowl. In addition, eight ALVE loci were identified. ALVE has not been found in any junglefowl species other than the red junglefowl, which supports the theory that commercial and indigenous chickens were domesticated from red junglefowl [31]. Although our phylogenetic tree showed separate clusters for gray junglefowl and chickens, these shared 12 ERV loci, one of which was not observed in red junglefowl. Genome analyses of junglefowl suggest that not only red junglefowl but also gray junglefowl contribute to the genetic diversity of poultry; hence, some ERV loci may have been inherited from junglefowl species other than red junglefowl. Although this analysis is based on a limited number of individuals and may not reflect all ERV loci within the population, analyzing more individuals in the future will allow us to accurately understand the ERV gene pool. The results of this study indicate that ERVs may act as powerful markers, similar to microsatellites and single nucleotide polymorphisms, for elucidating evolutionary relationships between breeds or species.

In the ERV-type analysis, multiple ERVs, including EAV-HP, EAV-51, and EAV-0, were identified with a particularly high frequency of EAV-HP insertions. On average, the contigs corresponding to ALVE were longer than those of the other ERVs, indicating diverse genomic structures. Many of the obtained contigs contained LTR regions, with some also including gag, pol, and env regions, which demonstrates the structural diversity of ERVs. Analysis of the 51 genes based on ERV insertion and gene expression revealed significant differences in the expression of *NAALAD2*, *GPC6*, and *PLSCR5*. For *NAALAD2* and *PLSCR5*, ERV insertion was associated with decreased expression, whereas *GPC6* showed increased expression. These results suggest that ERV insertions influence the functions of *NAALAD2* in neurotransmitter inactivation, *GPC6* in cancer cell migration and invasion, and *PLSCR5* in phospholipid scrambling of the cell membrane. The impact of ERV insertions may vary depending on the cell type, as specific LTR families are known to promote tissue-specific expression of host genes [32]. LTRs may function as alternative promoters when inserted near genes, with varying effects depending on the insertion site and sequence composition [33, 34]. Known cases of ERV effects in chickens include the blue eggshell color of Araucana, expression of immune- and inflammation-related genes in the liver, and feminization of plumage in Sebright breed males. Although ERVs are not confirmed to function as promoters in these instances, the significant changes in gene expression observed in this study suggest this possibility. Finally, while this study used RNA derived from skeletal muscle, further research is required to determine whether similar effects are observed using other cell types and to elucidate the specific mechanisms by which ERVs influence gene expression.

## Conclusion

We successfully identified the genomic loci of ERVs in broiler, Tosa-Jidori, and Yakido chickens. Some of these ERVs were inserted near genes and were correlated with the expression of *NAALAD2*, *GPC6*, and *PLSCR5* genes. Further research is required to determine whether these differences in expression are directly attributable to ERV insertion. Understanding the impact of ERVs on gene expression could reveal new insights into the genetic regulation and evolution of poultry. Future studies should aim to analyze a larger number of individuals and additional chickens to validate these findings and explore the broader implications of ERV insertions in genetic diversity and trait development.

## Materials and Methods

### Animal samples and genomic DNA and RNA purification

All experimental protocols and procedures were reviewed and approved by the Institutional Animal Care and Use Committee (IACUC) of Nippon Veterinary and Life Science University, Tokyo, Japan (Approval number: 2021-S34), ensuring compliance with ethical standards for animal research. The chickens selected for this study include Broilers, representing typical commercial chickens; Yakido, representing Malay-type native Japanese chickens; and Tosa-Jidori, representing traditional native Japanese chickens. Broilers were obtained from a farm in Kagoshima Prefecture. Tosa-Jidori chickens and Yakido chickens were sourced from the Livestock Research Center in Kochi Prefecture and Mie Prefecture, respectively. All chickens were euthanized via decapitation after isoflurane anesthesia at 35 days of age for broilers and 14 days for Tosa-Jidori and Yakido chickens. Blood and breast muscle samples were collected from chickens. Breast muscle was snap frozen in liquid nitrogen and stored at −80 °C until total RNA extraction. Genomic DNA was extracted from the blood samples using the QIAamp DNA Blood and Tissue Kit (Qiagen, Hilden, Germany), while DNA quality was evaluated via gel electrophoresis. Total RNA was extracted using an RNeasy Fibrous Tissue Mini Kit (Qiagen).

## Whole genome sequence (WGS) data

Genomic DNA from 16 individuals, including 8 broilers, 4 Tosa-Jidori, and 4 Yakido chickens was used to construct each 350-bp sequencing library with a TruSeq DNA PCR-Free Sample Prep Kit (Illumina Inc., San Diego, CA, USA). WGS was performed on 150-bp paired-end reads using an Illumina NovaSeq 6000 System (Illumina Inc.). Nucleotides with low-quality scores were trimmed from these reads, and adapters were removed with Trimmomatic v.0.36 [35] using the ILLUMINACLIP: TruSeq3-PE:2:30:10, LEADING:3, SLIDINGWINDOW:4:20, and MINLEN:30 settings. The reads were mapped to the *Gallus gallus* reference genome (GRCg6a, GenBank assembly accession: GCF_000002315.6) using the BWA-mem algorithm [23]. Data were generated with a Binary Alignment Map (BAM) format.

### Detection of non-reference ERVs

ERVs were detected as previously described [10, 36, 37]. The types of read pairs mapped to the reference genome were defined, and useful sequence reads were extracted. Most paired-end reads were obtained from a reference genome WGS map; however, mismatched read pairs also occurred with unexpected span sizes and orientations. Non-proper pairs were defined as those in which either the 5′ or 3′ end mapped to a contig sequence in the reference genome while the other mapped entirely or partially to an unexpected locus. Ends of a read pair that did not map to the reference genome were denoted as singletons, whereas unmapped read pair was used when both ends of a read pair did not map to the reference genome (Figure S4). Mismatched read pairs can provide insight into LTR-related loci as anchors; these were used with the RetroSeq software to detect non-reference transposon elements [24]. The process flow is illustrated in Figure S4. The ERV sequences used for RetroSeq were obtained from the National Center for Biotechnology Information (NCBI, Bethesda, MD, USA) and are listed in Table S5. The reference genome used was GRCg6a, containing only autosomes and sex chromosomes. In the RetroSeq “call” step, transposable element insertion positions (breakpoints) were estimated using reads detected in the “discover” phase, as previously reported [10, 36, 37]. The call step was set to ≥ 10 to reduce false positives, while the maximum read depth option per call was set to 10,000 to increase BAM coverage. All other RetroSeq options were used at their default values. A minimum of seven filter-level breakpoints was used, with breakpoints detected within 500 bp considered identical and excluded. The IGV [38] was used to detect loci containing TSDs; using the batch script functionality of IGV, a screenshot was obtained at each genomic locus detected using the RetroSeq analysis pipeline and carefully examined. The loci were presumed to be TSDs if they mapped to reads detected during the “discover” phase either from the 5′ or 3′ side, with an overlap ranging 1–10 bp. The 5′ and 3′ reads mapped within 150 bp of the TSDs were extracted using SAMtools [39]. The extracted read set was used to generate contig sequences using CAP3 software [40], which were subsequently used for a blastn search [41]. The lowest e-value was used to determine the ERV class. Each 200-bp sequence upstream and downstream of the breakpoint was extracted from the reference genome and subjected to blastn to eliminate the possibility of detecting ERV sequences in the reference genome. Subsequently, the loci that matched the ERVs were excluded from the analysis. Sequences that did not exist in the reference genome were deduced from the contiguous sequence within the region on either side of the TSD sequence or the 6-bp sequence adjacent to the 5′ and 3′ sides of the TSD when its length was insufficient.

### Analysis of Detected ERV Loci and Sequences

The detected loci in broiler, Tosa-Jidori, and Yakido chickens were compared with previously reported ERV loci in junglefowl [10]. The presence or absence of ERVs at each locus was represented as 1 or 0, respectively, to perform clustering among the species and subspecies. A resulting phylogenetic tree was constructed using the R software *ape* and *ade4* package [42, 43, 44] and “dist.binary” and “hclust” functions. Contigs containing ERV sequences at each locus were mapped to previously reported ERV sequences using the BWA-mem algorithm. Based on the mapping results, each contig was classified according to ERV species and further categorized by length. Homozygosity or heterozygosity was determined using IGV. Contigs were considered homozygous when reads mapped near the TSD were truncated. Conversely, when reads mapped across both the 5’ and 3’ ends of the TSD, the contig was considered heterozygous. The observed (Ho) and expected heterozygosities (He) for each chicken were estimated using the “popgen” function in the *PopGenKit* package [45] in R [42].

### Gene expression analysis using RNA-seq and relation

For RNA-seq analysis, the TruSeq cDNA library was constructed using the NEBNext® Ultra™ Directional RNA Library Prep Kit (Illumina, USA) after verifying the quantity and quality of the extracted RNA. Raw RNA-seq data were acquired using the Illumina NovaSeq 6000 platform, trimming was performed using the Trimmomatic software [35], and reads were mapped to the chicken reference genome (GRCg6a) using the STAR program [46]. Expression analysis was conducted using the RSEM program [47] to estimate TPM values. The ERV loci were identified and examined for insertions within the genes using IGV. For loci with ERV insertions within genes, TPM values were compared with those of individuals without ERV insertions at the same loci. A t-test was performed for each locus, with p-values adjusted using the Benjamini-Hochberg method, setting a threshold of statistical significance at α=0.05.

## Ethics approval and consent to participate

### Consent for publication

Not applicable.

### Data availability

The data used in this study are available from DDBJ (https://ddbj.nig.ac.jp/, BioProject accession: PRJDB17938).

### Funding

This work was supported by the Japan Society for the Promotion of Science Grant-in-Aid for Early-Career Scientists, Grant Number 22K14907.

## Acknowledgements

Computations were partially performed on the NIG supercomputer at the ROIS National Institute of Genetics.

## Authors’ contributions

S.I. designed the study and the main conceptual ideas, collected and analyzed the data, and wrote the manuscript. J.S., S.S., and D.I. collected the resources and data, supported, and supervised the writing of the manuscript. All authors discussed the results and commented on the manuscript.

