## Supplementary figures and images for "Endogenous Retrovirus Loci and Induced Changes in Gene Expression in Japanese Indigenous Chickens"

### Figure S1

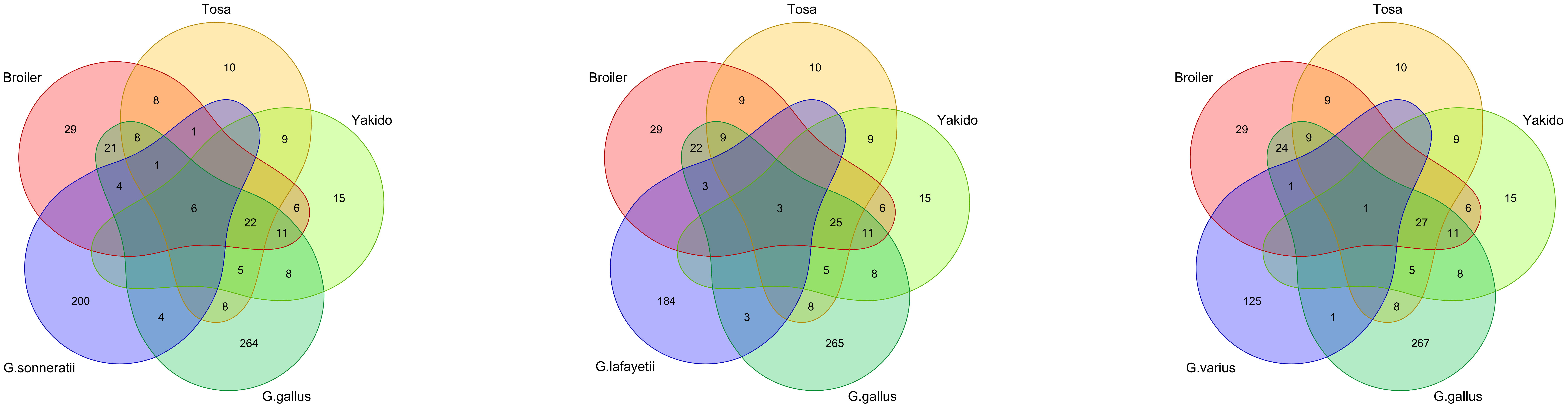

### Figure S2

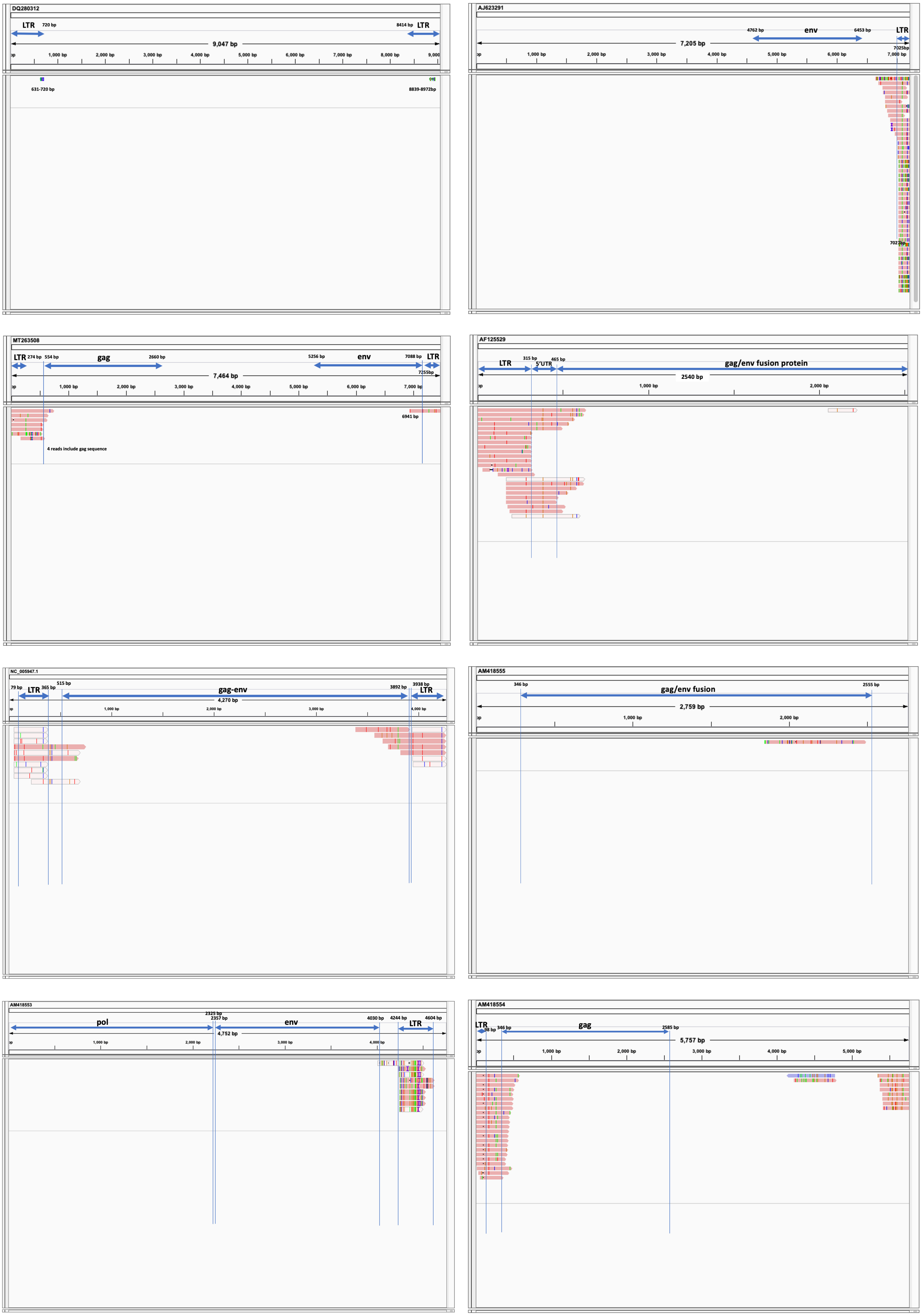

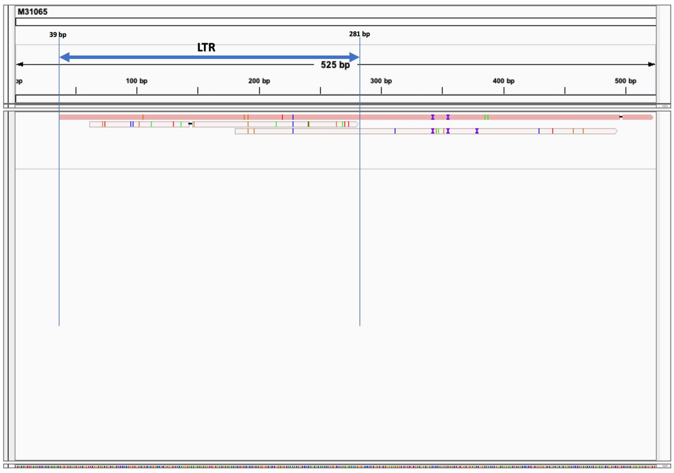

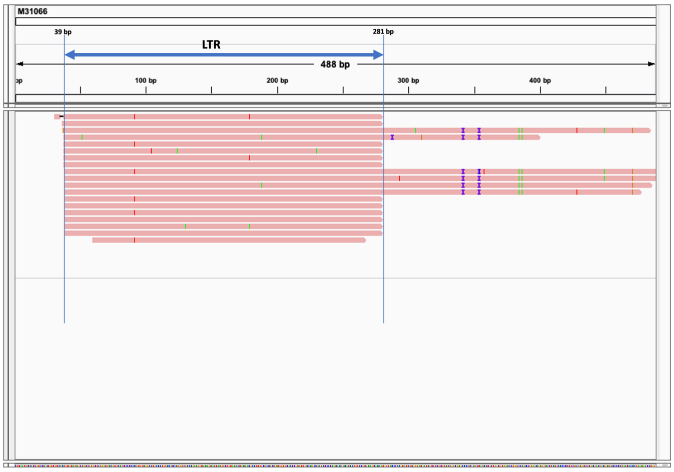

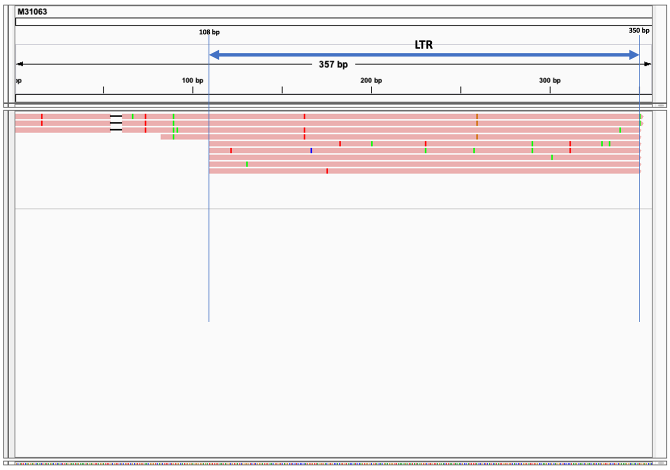

### Figure S3

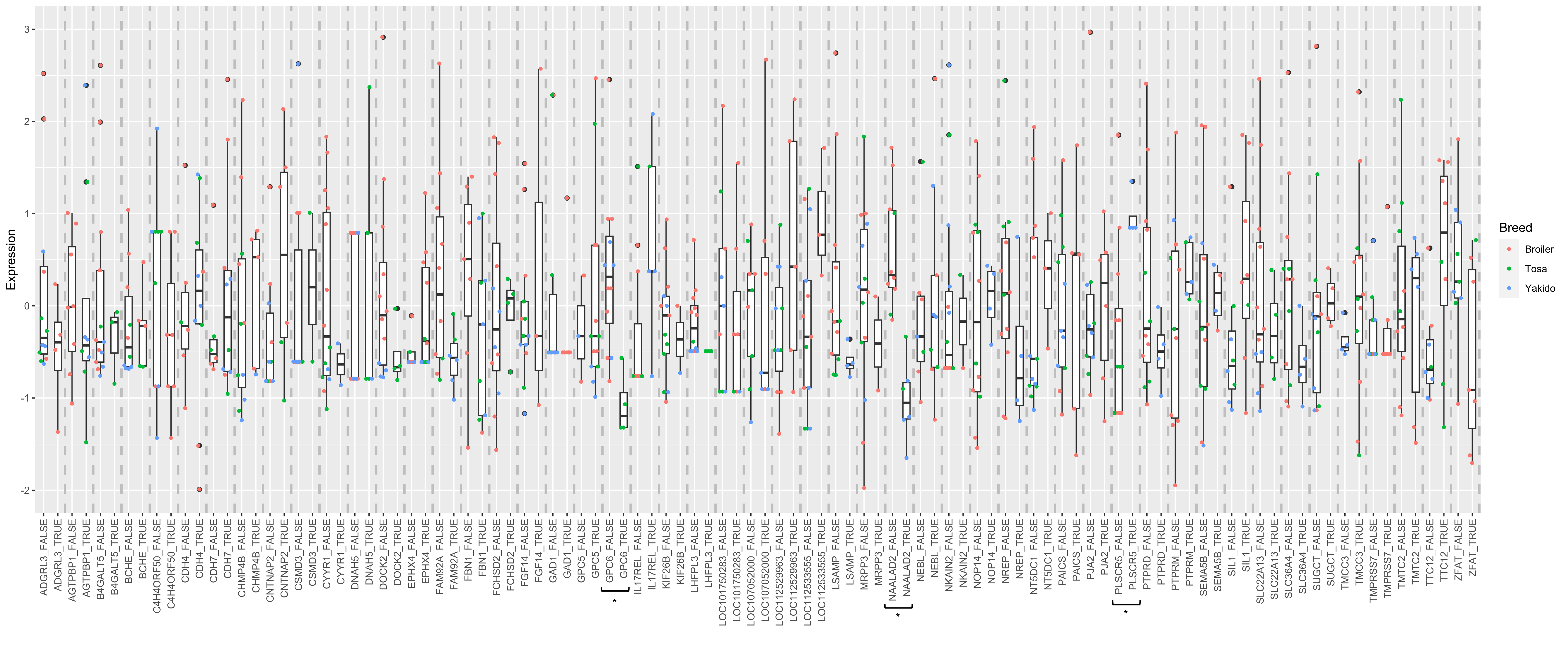

### Figure S4

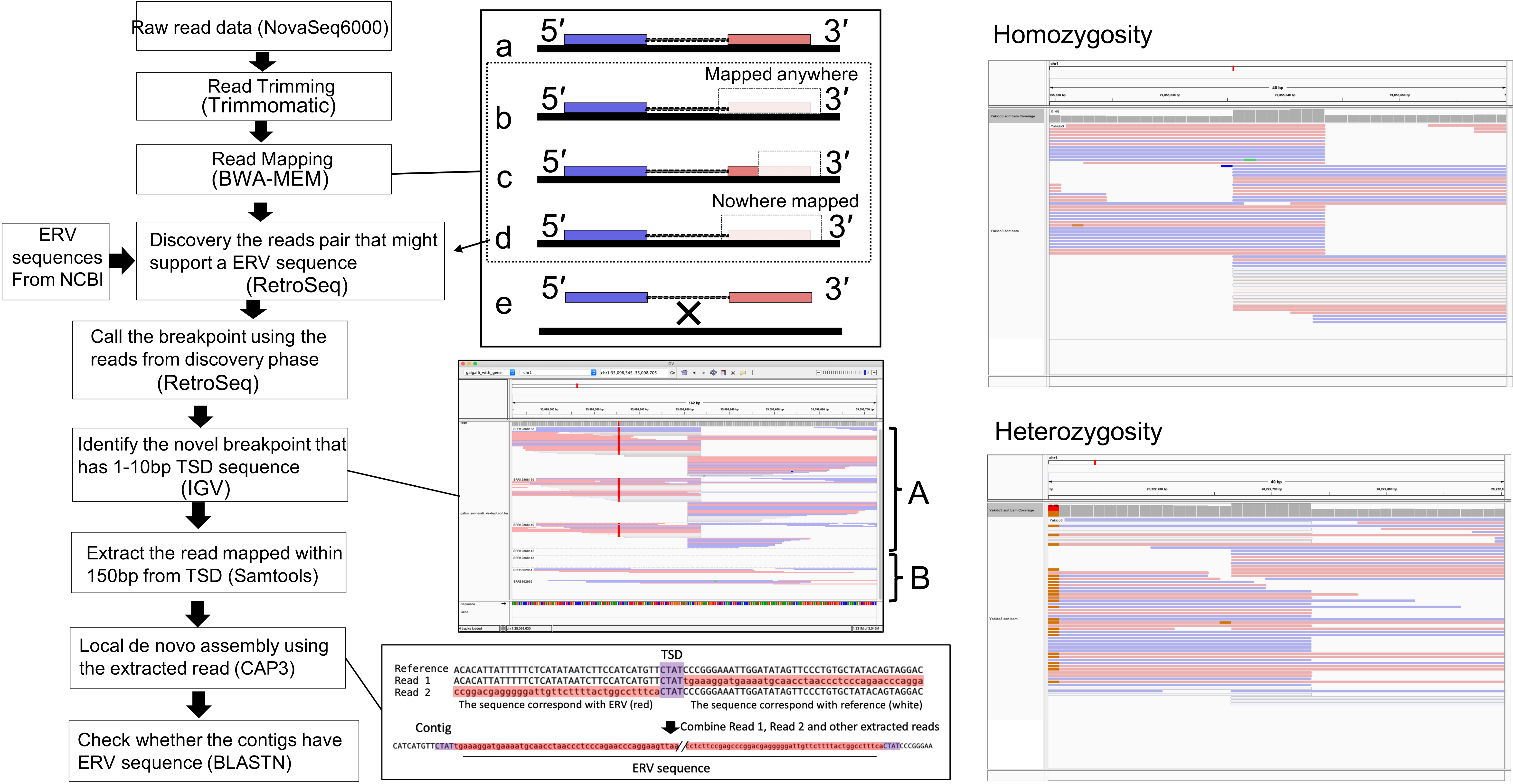
